# Stable and Oscillatory Hypoxia Differentially Regulate Invasibility of Breast Cancer Associated Fibroblasts

**DOI:** 10.1101/2024.03.26.586706

**Authors:** Wenqiang Du, Ashkan Novin, Yamin Liu, Junaid Afzal, Shaofei Liu, Yasir Suhail, Kshitiz

## Abstract

As local regions in the tumor outstrip their oxygen supply, hypoxia can develop, affecting not only the cancer cells, but also other cells in the microenvironment, including cancer associated fibroblasts (CAFs). Hypoxia is also not necessarily stable over time, and can fluctuate or oscillate. Hypoxia Inducible Factor-1 is the master regulator of cellular response to hypoxia, and can also exhibit oscillations in its activity. To understand how stable, and fluctuating hypoxia influence breast CAFs, we measured changes in gene expression in CAFs in normoxia, hypoxia, and oscillatory hypoxia, as well as measured change in their capacity to resist, or assist breast cancer invasion. We show that hypoxia has a profound effect on breast CAFs causing activation of key pathways associated with fibroblast activation, but reduce myofibroblast activation and traction force generation. We also found that oscillatory hypoxia, while expectedly resulted in a “sub-hypoxic” response in gene expression, it resulted in specific activation of pathways associated with actin polymerization and actomyosin maturation. Using traction force microscopy, and a nanopatterned stromal invasion assay, we show that oscillatory hypoxia increases contractile force generation vs stable hypoxia, and increases heterogeneity in force generation response, while also additively enhancing invasibility of CAFs to MDA-MB-231 invasion. Our data show that stable and unstable hypoxia can regulate many mechnobiological characteristics of CAFs, and can contribute to transformation of CAFs to assist cancer dissemination and onset of metastasis.

## Introduction

Breast tumor growth commonly outruns its supply of oxygen, creating localized hypoxia^1^. Hypoxia can act as a key microenvironmental cue in the cancers, regulating many aspects of cancer progression and its transformation to a malignant disease^2^. The primary cellular regulator of hypoxia is a transcription factor (TF), Hypoxia-inducible factor-1 (HIF-1), which acutely responds to lack of molecular oxygen. HIF-1 is composed of two subunits, with HIF-1a being the regulated subunit, which is stabilized in lack of oxygen. Upon stabilization, HIF-1a binds to constitutively expressed HIF-1b and transcriptionally regulates a large number of downstream genes by acting on the HIF-responsive element (HRE) in the cis-regulatory region of target genes^3^. However, in recent years with the advent of molecular imaging, there is an increased appreciation of the spatiotemporal heterogeneity in hypoxia presence in the tumor microenvironment. Although hypoxic regions were known to be non-uniformly distributed, evidence is accumulating for fluctuations in oxygen concentrations in many regions of tumor^4-5^. Many mechanisms are attributed to these fluctuations, or oscillations, including leaky and immaturely developed vasculature, rapid consumption of oxygen by cells with not stabilized HIF-1, as well as fluctuations in temperature. We have shown that oscillatory hypoxia can specifically upregulate gene expression which are highly enriched in human cancers, and prognosticate low survival in breast cancer patients^6^. There are other reports on hypoxic oscillations resulting in malignant transformation of cancer^4^.

Although role of hypoxia has been well studied in cancers, its role in regulation of other cell types in the tumor microenvironment is relatively less studied^7^. As tumors grow, many other non-cancerous cells are recruited into the tumor niche, including the stromal and immune cells^8^. Stromal cancer associated fibroblasts (CAFs) are now considered to contribute significantly to cancer dissemination^9-10^. We have shown that stromal resistance, or abetment, to cancer invasion is a selected and regulated phenotype, identifying many genetic and non-genetic factors which contribute to it^11^. As epithelial tumors grow and break open the basal lamina, stromal fibroblasts get activated in a classic wound healing response^8^. This involves their transformation into myofibroblasts, characterized by increased contractile force generation, and change in matrix production^12^. However, our understanding of how hypoxia, and fluctuations in hypoxia, influence myofibroblast activation, as well as stromal invasion is limited. As HIF-1 is ubiquitously expressed in all cell types, hypoxia is likely to have an effect on CAFs (**Figure 1A**). To directly identify how hypoxia, and oscillatory hypoxia regulate gene expression in CAFs, we isolated fibroblasts from human breast cancer biopsy, and performed RNAseq after conditioning the cells with hypoxia, and a regiment of intermittent hypoxia. We found that hypoxia expectedly changed key biomechanical pathways, but surprisingly oscillatory hypoxia resulted in expression of many genes associated with mechanical activation. Using traction force microscopy of CAFs in hypoxia, and oscillatory hypoxia, we found that oscillation increased the heterogeneity in force generation, generating a small subpopulation of CAFs with significantly high traction force not evident in normoxia and hypoxia. Finally, using a nanopatterned stromal invasion assay, we demonstrate that oscillatory hypoxia can significantly enhance the stromal vulnerability to breast cancer invasion, increasing the deep dissemination of cancer cells into the stromal compartment. Overall, our work highlight that oscillations in tumor hypoxia has a direct effect on non-cancer CAFs, transforming their mechanical phenotype, and contributing to the malignant transformation of breast cancer.

**Figure 1.**
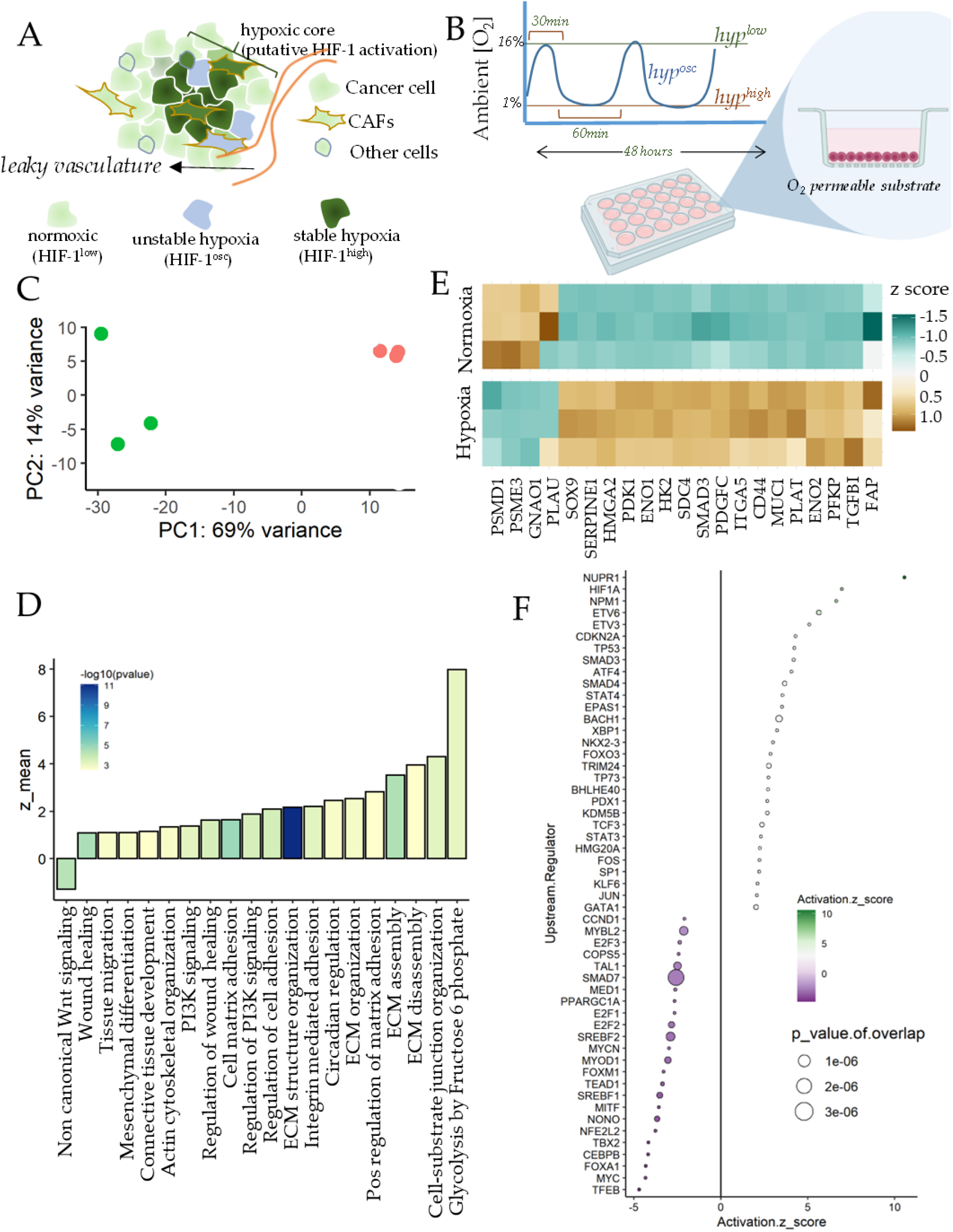
Hypoxia results in fibroblast activation in breast CAFs. (A) Schematic showing a hypoxic tumor locality with regions of stable and fluctuating (or oscillatory) hypoxia sensed by both cancer, and non-cancerous cells including cancer associated fibroblasts (CAFs). (B) Schematic showing the experimental plan. For all experiments, cells were cultured on an O2 permeable surface as O2 diffusion is extremely low in aqueous medium. Ambient O2 levels were maintained either as normoxia (16% O_2_), hypoxia (1% O_2_), or oscillatory (1 hour of hypoxia followed by 30 min of normoxia for 48 hours). (C) Principal Component Analysis (PCA) of gene expression in CAFs isolated from breast treated with normoxia (red circles) or hypoxia (green circles); each circle represents an independent biological sample. (D) Mean z-scores of top gene ontologies activated by hypoxia vs normoxia in breast CAFs. (E) Heatmap showing z-score of expression for selected genes contributing maximally to the activation score of GOs in D. (F) IPA predicted top Upstream Transcriptional Regulators for differentially expressed genes in hypoxia; size of the bubble represent p-value, while x-axis and color map show predicted z-score of regulator activation.

## Results

### Hypoxia induces a pro-fibrotic gene expression outcome in Breast Cancer Associated Fibroblasts (CAFs)

To assess the effect of hypoxia on CAFs, we first isolated fibroblasts from breast cancer tissue biopsy from a patient with ER+PR+HER2-breast cancer, and treated a monolayer of CAFs with 1% O_2_ for 48 hours. Molecular O_2_ has extremely low solubility in aqueous solutions like tissue culture media^13^, and can take many hours to equilibrate from ambient O_2_ atmosphere. Because we also aimed at observing the effect of oscillations in hypoxia, we cultured the cells on O2 permeable silicone surface, so that ambient O_2_ is immediately available to the cells from the bottom of the plate (**Figure 1B**). RNA was isolated immediately after hypoxic conditioning, sequenced, and analyzed for differential expression of genes between normoxia and hypoxia.

Principal Component Analysis showed a large separation in gene expression for normoxia and hypoxia samples (**Figure 1C**). To assess the broad contours of gene expression changes, we identified the key gene ontologies (GOs) activated by hypoxia. We found many GOs significantly activated in hypoxia were related to regulation of extracellular matrix assembly, and organization, actin cytoskeletal reorganization, tissue migration, and wound healing (**Figure 1D**). In addition, we also found activation of glycolytic processes and circadian regulation, which are well known to be regulated by HIF-1^14^. Other notable GOs were those associated with cell-matrix adhesion, including integrin activation, cell-matrix adhesion, and cell-substrate junction organization. Crucially, non-canonical Wnt signaling was downregulated by hypoxia in breast CAFs. Overall, hypoxia appeared to directly regulate many matricellular and biomechanical pathways associated with wound healing, and stromal response to cancer dissemination (**Figure 1D**). We selected the genes prominently contributing to activation of these pathways from the top activating sets, finding key genes responsible for fibroblast activation, wound healing, and matrix interaction, as well as those related to glycolysis (**Figure 1E**). These included genes encoding key components of the Transforming Growth Factor beta (TGFβ) pathway, involved in myofibroblast transformation of fibroblasts, including transcriptional co-regulator Smad3, TGFβ-1 ligand itself, and TGFBI encoding the TGFβ-inducible protein. In addition, we found other key genes associated with CAFs interaction with matrix to be increased by hypoxia, including FAP encoding Fibroblast-activating Protein, SDC4 encoding Syndecan-4, MUC1 encoding mucin-1, HMGA2 which is an independent marker for stromal activation^15^, ITGA5 encoding Integrin a5 subunit responsible for enhanced tumor-stroma interaction, and CD44, a membrane glycoprotein involved in ECM regulation of TGFβ pathway. While hypoxia increased expression of several genes regulating CAF-matrix interaction, those downregulating matrix degradation were also increased, including SERPINE1, PLAT, and PLAU, key inhibitors of the matrix metalloproteinases. Furthermore, SOX9, which is a key regulator of fibroblast activation was also increased in hypoxia, as well as PDGFC^16-17^. In addition to the genes associated with fibroblast activation, other key genes were involved in the glycolytic shift in metabolism, including PFKP encoding a member of the phosphofructosekinase family, as well as ENO1 and ENO2 encoding enolases, HK2 encoding hexokinase-2, and PGK1 encoding phosphoglyceratekinase. Hypoxia also resulted in reduced expression in FZD1 encoding Frizzled-1, SMURF2 which is a Smad specific ubiquitin ligase and is an inhibitor of fibrosis (**Figure 1E**).

Upstream TFs predicted to be regulating the differentially expressed genes in hypoxia included HIF1A, TP53, EPAS1 which encodes for Hif-2a, as well as Smads, which regulate TGFb signaling (**Figure 1F**). In addition, we also found other nuclear regulators like Nucleophosmin (NPM1), and Nuclear Protein 1 (NUPR1), as well as XBP1 which regulates unfolded protein folding response. Hypoxia induced gene expression also indicated inhibition in activity of Smad7, which acts as an inhibitory TF in TGFb pathway, SREBF1, and SREBF2 encoding for sterol regulatory element binding TF 1, and 2, as well as MYOD1 encoding for myogenic differentiation factor 1. Overall, transcriptomic analysis strongly suggested that hypoxia resulted in increased fibroblast activation in breast CAFs, which was surprisingly not accompanied by myofibroblast transformation, evidenced by reduced MYOD activity^18-19^.

### Hypoxia upregulates genes associated with actomyosin transformation of CAFs

To further explore the specific genes and pathways associated with hypoxia induced transformation of CAFs, we performed Gene Set Enrichment Analysis (GSEA) for the differentially expressed genes (DEGs) between hypoxia and normoxia in CAFs. GSEA is a non-parametric statistical test to assess if a given set of DEGs are preferentially enriched in a given functionally meaningful gene-set vs in random sets, and normalized to the size of the gene-set^20^. GSEA analysis confirmed our initial finding that hypoxia can induce a broadly pro-fibroblast activation program in CAFs, typically mediated by activation of TGFβ signaling, and characterized by increased production of extracellular matrix, and reorganization of cytoskeletal organization. We found key gene ontologies to be significantly enriched by DEGs in hypoxia including cell-matrix adhesion, connective tissue development, and ECM assembly with high normalized enrichment scores (NES). Leading edge genes of these pathways revealed several key genes associated with the fibroblast transformation of CAFs in hypoxia.

Many genes encoding ligands and receptors of TGFβ signaling including ACVR1C, ACVR2B, BMP6, BMP7, NODAL, as well as SMAD3, and SMAD7 which are the endpoint transcriptional regulators of TGFβ signaling. In addition, surprisingly we also found many genes encoding the various subunits of inhibins, including INHBA/B/C/E in hypoxia. Typically inhibins and activins can act oppositely to regulate TGFβ signaling, but many inhibin subunits interact with both inhibins and activins^18, 20^. Hypoxia therefore seem to not just activate TGFβ signaling, but also its regulators, including the negative ones, suggesting a more nuanced regulation of myofibroblast activation (**Figure 2A**). This was also observed in the upregulation of GREM1, which encodes Gremlin-1, a secreted ligand negatively regulating TGFβ activation (Figure 2B). As mentioned, both ECM assembly, and cell matrix adhesion were enriched by DEGs in hypoxia (**Figure 2B-C**). Leading edge genes included ECM molecules including those encoding for fibronectin (FN1), and collagens (COL13A1, COL1A2), Laminin (LAMB2, and LAMB4), Nidogen (NID2), as well as hyaluronon (HAS1, HAS3) as well as CD44, which can act as a receptor for hyaluronans. Interestingly, we also found increased expression for PKD1 which encodes polycystin-1, a transmembrane Ca2+ ion channel, as well as SLC9A1 encoding a Na+/H+ exchanger. pH dysregulation can alter cancer manifestation, and more and more evidence are building on pro-tumor functionalization of CAFs by pH^21-22^.

**Figure 2.**
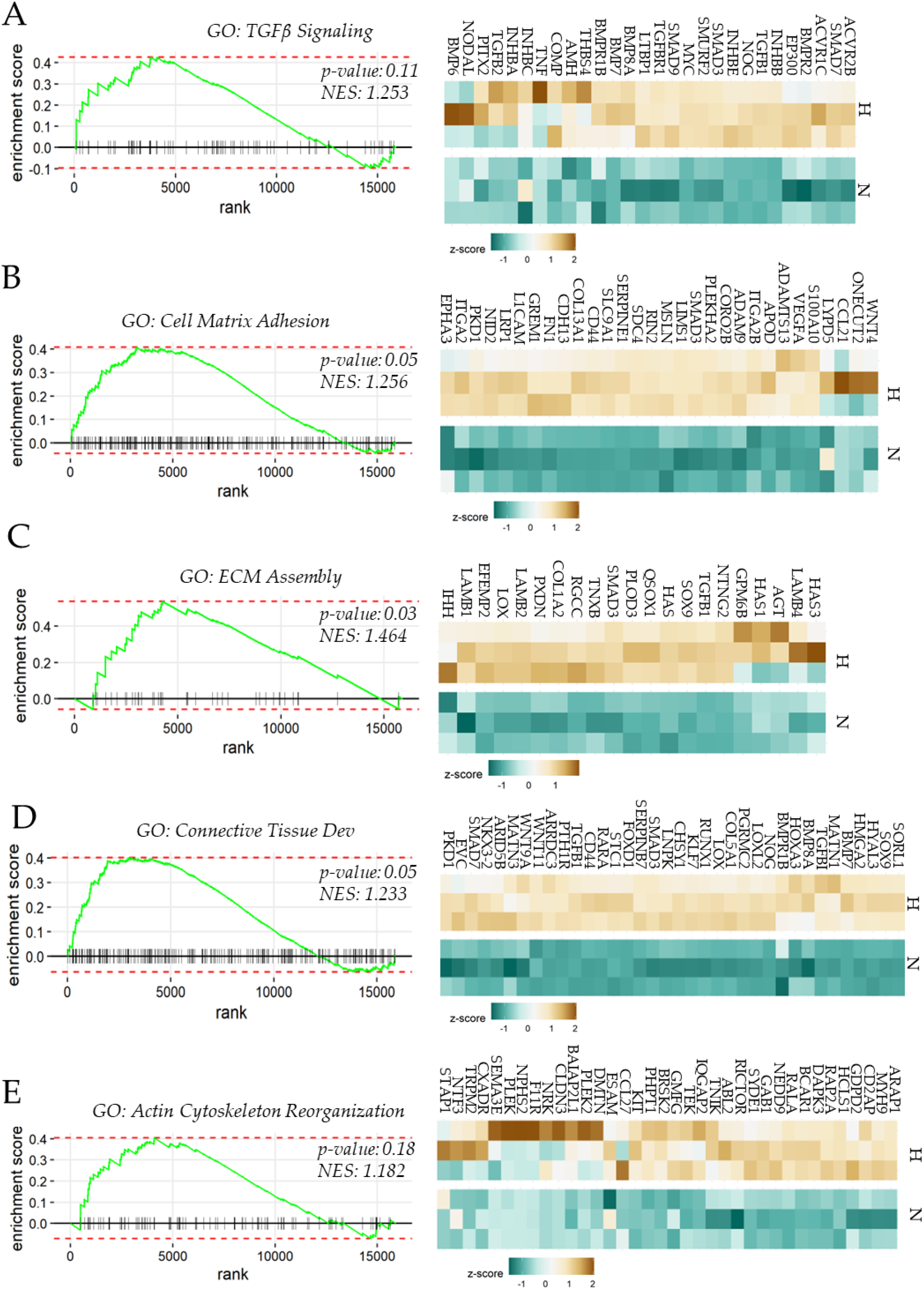
Gene Set Enrichment Analysis show hypoxia increases gene expression associated with matrix production and adhesion in breast CAFs. GSEA plots (left), and heatmap of leading edge genes (right) for differentially expressed genes in hypoxia vs normoxia in (A) TGFb signaling, (B) Cell Matrix Adhesion, (C) Extracellular Matrix Assembly, (D) Connective Tissue Development, and (E) Actin Cytoskeleton Reorganization. p-value and enrichment scores are provided for each GO, and genes are ranked in order of their fold change between hypoxia and normoxia. H: Hypoxia, N: Normoxia. Scalebar for z-scores shown in each heatmap.

**Figure 3.**
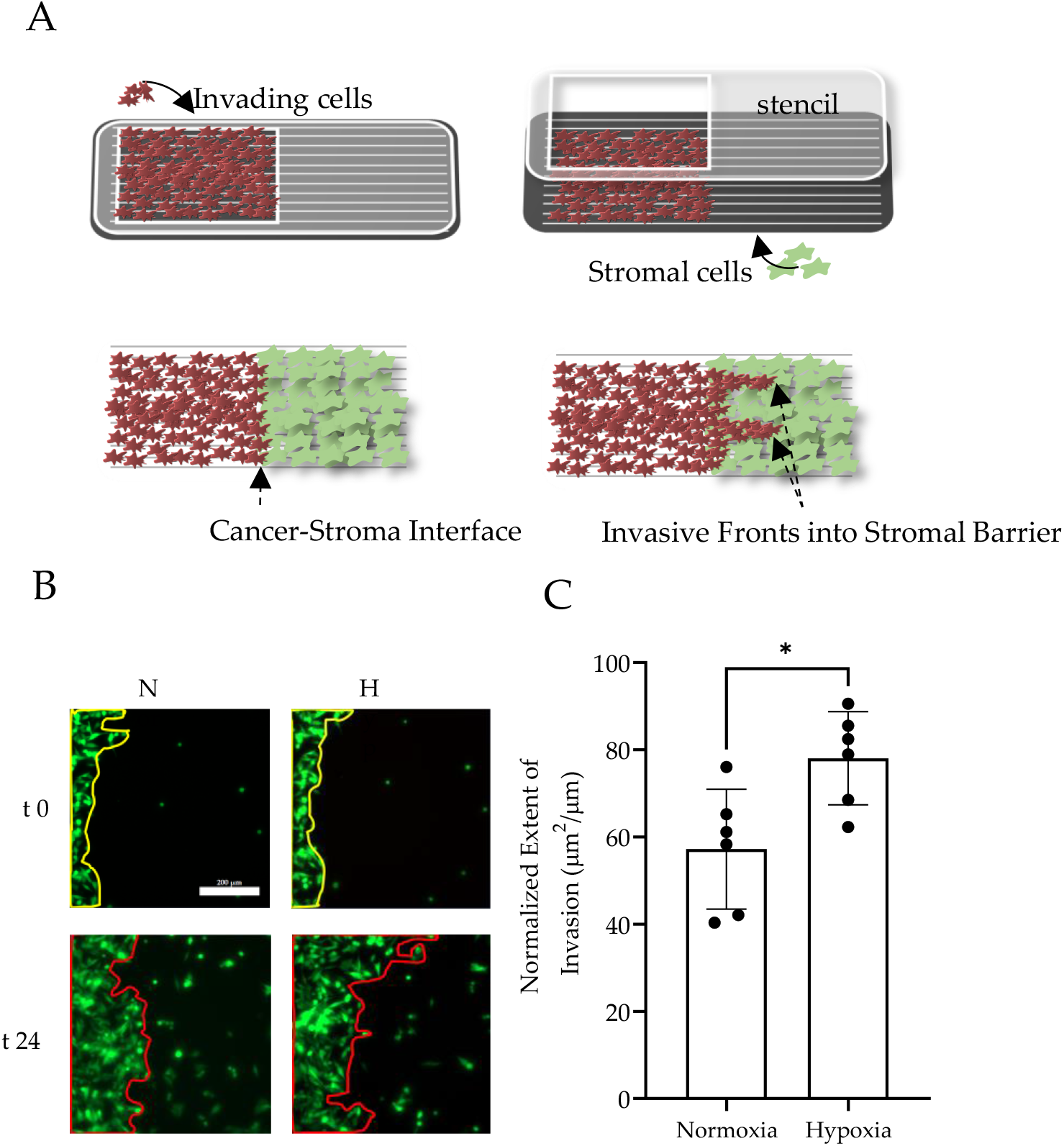
Hypoxia increases invasibility of breast CAFs. **(A)** Schematic showing ANSIA (Accelerated Nanopatterned Stromal Invasion Assay). Fluorescently labeled MDA-MB-231 cells, and unlabeled breast CAFs are patterned as confluent monolayers using a polymeric stencil on an anisotropic nanopatterned surface with the underlying grooves aligned orthogonal to the cancer-stromal interface. Fluorescence microscopy is used to measure extent of cancer invasion into the stromal compartment. (B) Representative images of MDA-MB-231 (green) monolayer interfaced with breast CAFs (black region, right) at 0, and 24 hours of experiment in normoxia (N), and hypoxia (H). (C) Extent of MDA-MB-231 invasion into the stromal compartment normalized to the interface length; n = 6 interface locations. *: p-value < 0.05 determined by Student’s-t-test; Error bars: stdev.

Leading edge genes revealed in GSEA enrichment of “connective tissue development” (**Figure 2D**) included SOX9, a key developmental TF which has been implicated in fibroblast activation and fibrosis^23^, KLF7 a TF involved in osteoclast differentiation^24^ and has been implicated in cancers^25-26^, HMGA2 which is induced by Fibroblast Growth Factor 2 and is therefore associated with fibroblast activation^27^, and has been reported to be an independent stromal marker for pancreatic adenocarcinoma. In addition to the gene ontologies associated with wound healing response and fibroblast activation, we also found that hypoxia resulted in enrichment of actin cytoskeletal reorganization. Surprisingly, among the leading edge genes, we did not find many myosin chain encoding genes except MYH9 which increased in hypoxia (Figure 2E). Other notable genes were ARAP1 which encodes an adapter protein with several signaling docking domains including PH1, RhoGAP, and Ankyrin repeats, and has been considered as a signaling hub to converge Rho and Arf signaling^28^. RAP2A and RALA encoding members of the RAS oncogene family, BCAR1 which encodes for Breast Cancer Anti-Estrogen Resistance Protein 1, and KIT, a member of the protooncogene KIT family. Interestingly, we also found increase in CLDN3 which encodes for Claudin-3, a tight junction protein, which could contribute to the collective migration of fibroblasts^29^.

### Hypoxia increases stromal invasibility of breast CAFs

CAFs are now considered as key mediators of cancer dissemination. However, the debate is yet not settled whether CAFs universally promote, or resist cancer invasion. Fibroblast activation is a physiological response to the wounding caused by basal lamina disruption by a growing tumor, and is aimed at containing and resolving the wound. Indeed, stromal invasibility to cancer invasion is a regulated and selected phenomenon, which we have shown is a secondary outcome of the differential control of placentation during pregnancy. At some stage during the cancer progression, fibroblasts are transformed to be assistive in cancer invasion. We have shown in pancreatic ductal cell carcinoma, that this transition is attained at the onset of metastasis, wherein the CAFs transform from a homogeneous population with a resistive potential to a heterogeneous stromal population which either does not resist cancer invasion, or actively promote it. Here, we asked how does hypoxia, which is a common microenvironmental variable in tumors, regulate stromal invasibility in breast cancers.

Stromal invasion by cancer, unlike cancer migration, is a slow process to measure experimentally. We, therefore, have developed an assay to measure stromal invasion in an accelerated manner by using an underlying anisotropic nanopatterned matrix to align cellular actomyosin assemblage in the same direction. The assay, called ANSIA (Accelerated Nanopatterned Stromal Invasion Assay) allows live analysis of the phenomenon of stromal invasion at the interface of monolayers of stromal fibroblasts and fluorescently labeled cancer cells, with the interface arranged orthogonal to the underlying nanofibers using stencil-based cell patterning. Using ANSIA, we measured breast CAF’s resistance to MDA-MB-231 invasion in normoxia and hypoxia.

To quantitatively measure the stromal invasion in hypoxia, we measured the difference between the spatial positioning of the initial CAF-cancer interface, and followed the changes in the interface for 24 hours. The difference in the aerial advancement of MDA-MB-231 cells was normalized to the initial length of interface. The normalized extent of invasion measured showed that hypoxia could significantly increase stromal invasion, suggesting that the gene expression changes induced by hypoxia reduce stromal resistance to cancer invasion.

### Oscillatory hypoxia both reverses hypoxia induced gene expression in CAFs, while enhancing actin network formation

As hypoxia had a large effect on stromal invasibility, we asked if fluctuations in hypoxia could mitigate the hypoxia-induced vulnerability to cancer invasion. As oxygen availability is intermittent in many intra-tumor regions, it results in fluctuations in hypoxia. Oscillatory hypoxia could be construed as an intermediate biological signal, averaging the normoxic and hypoxic response of the cell. We had previously found that emergent oscillations in HIF-1 activity can allow cancer cells to escape hypoxia driven inhibition of cell proliferation. This phenomenon suggested an intermediate response to hypoxia and normoxia. However, we had surprisingly found that many genes responded quite differently to oscillatory hypoxia, sometimes in the opposite direction to stable hypoxia, suggesting that gene expression by oscillatory hypoxia is not trivially predicted to be intermediate in between hypoxia and normoxia. Although oscillatory hypoxia has been studied in cancer cells, its effect on CAFs has not yet been elucidated.

We therefore subjected CAFs to an oscillatory or intermittent hypoxic dose using a dynamic tri-gas controller on a tissue culture plate with an O2 permeable bottom surface (**Figure 4A**). Cells were isolated immediately after the experiment during an oxidation cycle, RNA isolated and sequenced. A broad analysis suggested that the most significant changes between oscillatory and stable hypoxia were related to reversal in the HIF-1 induced glycolytic metabolism. Kegg signaling pathway analysis highlighted TCA cycle, glutathione metabolism, oxidative phosphorylation, as well as fatty acid metabolism (**Figure 4B**). However, we also found that many of the GOs which had increased in hypoxia showed a reversal in oscillatory hypoxia (**Figure 4B**). These included TGFb signaling, and focal adhesion, as well as ECM-Receptor interaction. These results were in accordance to the default expectations that oscillatory input should average the high and low inputs, with an intermediate “sub-hypoxic” downstream response.

**Figure 4.**
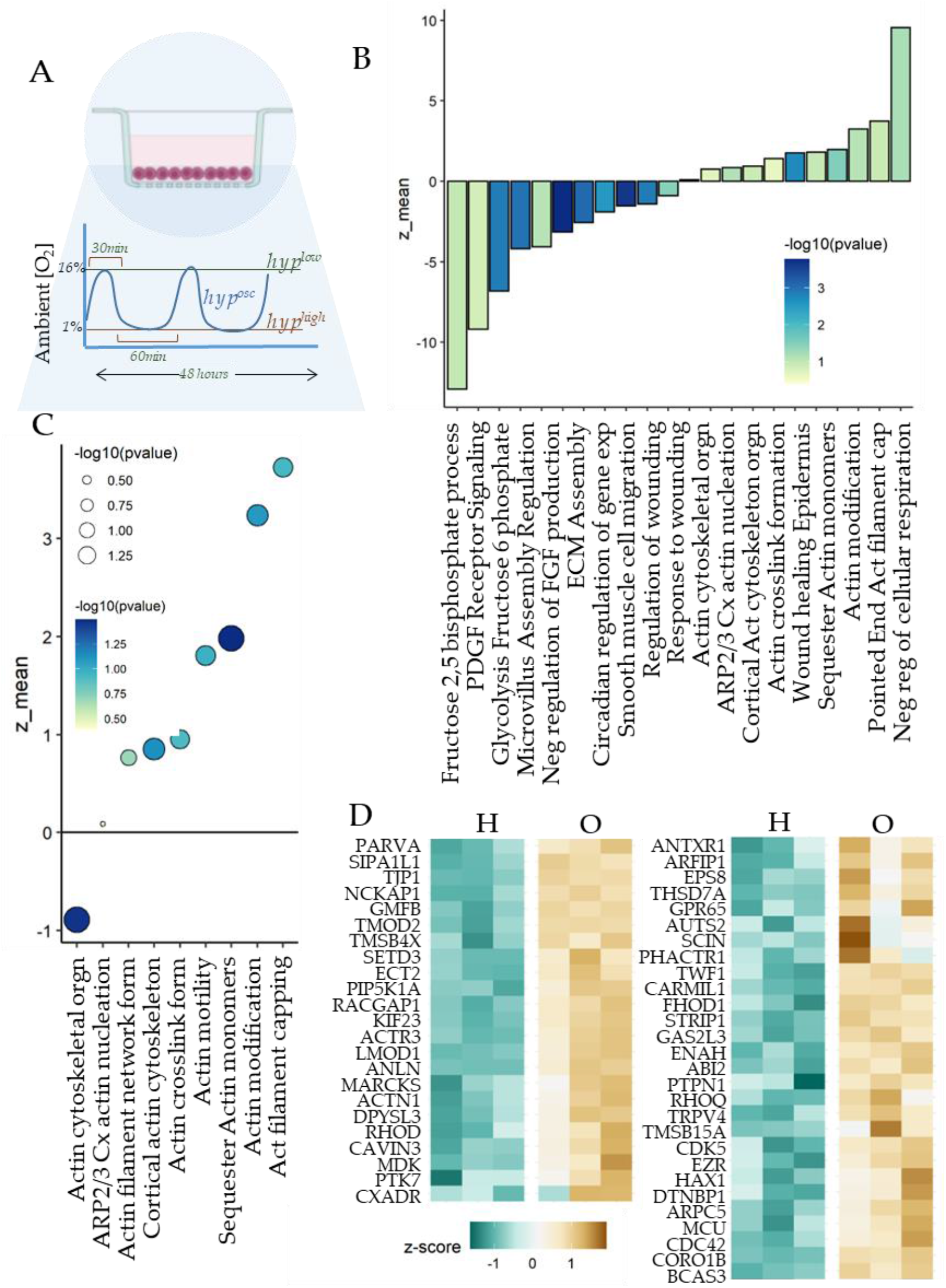
Oscillatory hypoxia increases genes associated with actin polymerization in breast CAFs. **(A)** Schematic showing the experimental setup with breast CAFs cultured in multi-well plates with O_2_ permeable bottom surface to allow rapid bioavailability of ambient O2 to the cultured cells; also shown is the experimental protocol. (B) Activation scores for gene ontologies (GO) activated by oscillatory hypoxia vs stable hypoxia; Color in the bar show p-value. (C) Bubble plot showing z-score for GOs which were activated in oscillatory hypoxia even more than in stable hypoxia. (D) Heatmap showing genes contributing maximally to the selected GOs in C.

However, GSEA analysis of GOs also showed specific ontologies which were activated in oscillatory hypoxia (vs stable hypoxia) related to actin network formation in cells (**Figure 4C**).

These included gene-sets related to actin modification, actin cross-link formation, cortical cytoskeletal organization, and regulation of Arp2/3 based complex. The leading edge genes of these pathways also highlighted genes which had an “oscillatory specific” effect on expression, suggesting that oscillatory hypoxia caused changes in CAF actin cytoskeletal organization differently from hypoxia (**Figure 4D**). These genes included LMOD1 encoding Leiomodin1 and TMOD1/2/3 encoding tropomodulin isoforms, both regulating myosin contractility, several actin binding proteins including TWF1 encoding twinfilin, PARVA encoding Parvin, and ANLN encoding anilin. In addition, we found many other actin capping proteins encoded by genes including CORO1B encoding Coronin-1B, ARPC5 and ARPC3 encoding Arp2/3 subunit. Overall, our data showed that while oscillatory hypoxia reduced TGFb activation and actomyosin maturation, it also resulted in the increase of actin crosslink formation and maturation. We therefore sought to investigate the phenotypic effect of these dichotomous changes.

### Fluctuations in hypoxia increases heterogeneity in CAF mechanics enhancing stromal invasibility

Our data showed that while hypoxia caused fibroblast activation in breast CAFs, accompanied by upregulation of TGFb signaling, increased matrix production, and integrin mediated matrix adhesion. Oscillations in hypoxia resulted in partial reversal of these pathways expectedly, but also surprisingly caused increased activation in actin crosslinking and maturation. These data suggested that oscillatory hypoxia may be differently sensed by CAFs than stable hypoxia in terms of their actomyosin crosslinking, likely contributing to increase in contractile machinery maturation. Furthermore, considering that while TGFb signaling was reversed, and actin crosslinking was increased, oscillation may produce increased heterogeneity in the mechanical response in CAFs.

We therefore functionally tested the effect of stable and oscillatory hypoxia on contractile force generation in CAFs using Traction Force Microscopy (TFM). Using fluorescent beads embedded in a labile hydrogel with a known Young’s modulus, which served as the substrate for CAFs, we measured the distribution of force generation at subcellular resolution after subjecting CAFs to 48 hours of normoxia, hypoxia, or oscillatory hypoxia (**Figure 5A**). As cells form matrix-integrin adhesion bonds, which is coupled to the intracellular actomyosin assembly, the contractile force generation is transmitted to the substrate, which could be measured by the displacement of fluorescent beads using microscopy.

**Figure 5.**
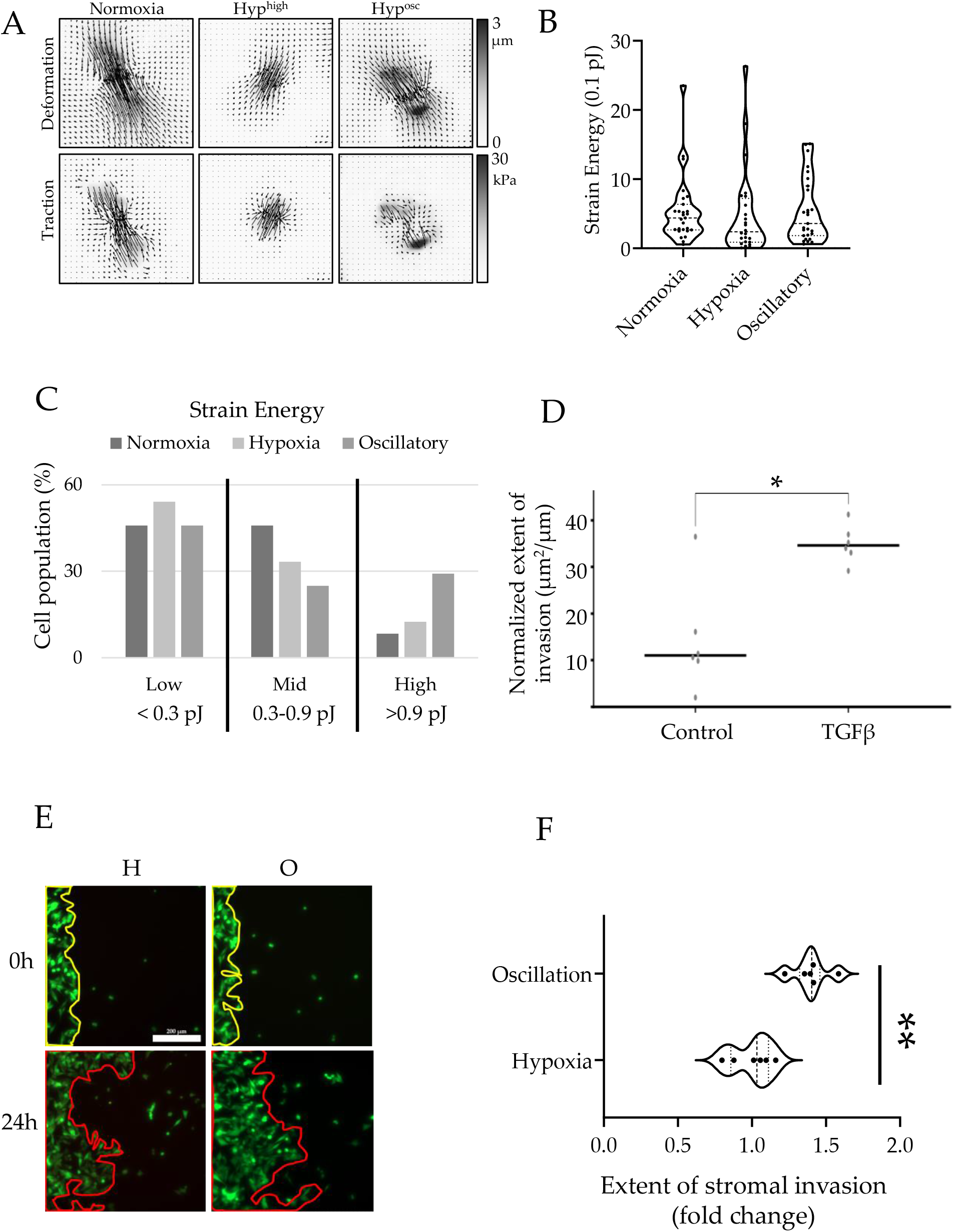
Oscillatory hypoxia increases stromal invasibility of breast CAFs. **(A)** Traction force microscopy derived vector maps of representative cells in normoxia, stable hypoxia, and oscillatory hypoxia showing deformation of the underlying hydrogel, and computed traction forces, with quantification shown in (B); no statistical significance was observed when the means were compared across the three conditions. (C) Composite histogram showing relative distribution of cells (n > 20 cells for each condition) binned at low, medium, and high force generation values. (D) Normalized extent of invasion of A375 cancer cells into the BJ compartment in ANSIA, when BJ were untreated, or previously treated with 10ng/ml TGF-β for 24 hours and seeded in the stromal compartment. (E) Representative images of MDA-MB-231 (green) cells invading into breast CAFs in stable hypoxia, and oscillatory hypoxia at 0h, and 24 hours of experimentation. (F) Fold change in the degree of invasibility of breast CAFs in oscillatory vs stable hypoxia; n = 6 locations; thick dashed line: median, thin dashed line: 10, and 90 percentile of distribution; **: p-value < 0.01 by Student’s-t-test.

When observed over 30 different cells, we found that although no statistical differences were found between the means for each condition (**Figure 5B**), observing the distribution of traction force generated per cell, we surprisingly found that oscillatory hypoxia increased the percentage of cells with high traction force generation (∼30%) (**Figure 5C**). Does stromal fibroblast contractility directly regulate their resistance to invasion from carcinomas? Transforming growth factor beta (TGFβ) is the primary cytokine involved in myogenic transformation of stroma observed in cancers. As CAFs are a heterogeneous group of cells, we treated BJ fibroblasts, a cell line derived from skin, with 25 ng/ml TGFβ for 24 hours and tested their potential to the relevant A375 melanoma invasion. We found that TGFb treatment significantly enhanced stromal invasibility (**Figure 5D**). We then asked if the stromal invasibility was altered in oscillatory hypoxia. ANSIA based stromal invasion assay showed that oscillatory hypoxia further decreased resistance of CAFs to invasion, or transformed CAFs towards a more invasible phenotype even more than stable hypoxia (**Figure 5E-F**).

Overall, our data show that oscillations in hypoxic stimuli could further enhance stromal invasibility, but likely through a different mechanism than hypoxia alone, by increasing the variability in CAF contractile force generation. Coupled with our previous observation that while hypoxia increased fibroblast activation programs, it also decreased TF activity associated with myofibroblast transformation, and that oscillation in hypoxia increased actin networking and maturation, our data suggests that regions with unstable hypoxia may increase CAF’s assistance to breast cancer dissemination.

## Discussion

Breast cancer is a leading cause of death in women, with a high medical, psychological, social, and financial cost. For long the focus on understanding progress of breast cancer from a primary tumor to a metastatic disease has remained on cancer cells themselves. However, in recent years, it has become increasingly evident that non-cancerous cells, particularly the fibroblasts and the immune cells in the microenvironment play active roles in transforming breast cancer into a malignant and lethal disease. Invasion into the stromal tissue is one of the early steps in the metastatic cascade. Although a large literature has built up about the characterization of stromal fibroblasts in the tumor microenvironment, our understanding about how the microenvironment itself influences fibroblasts is limited. Tumor microenvironment is highly specialized, and presents aphysiological concentration ranges of various biochemical and mechanical factors. Of particular consequence is the lack of oxygen, or hypoxia, which builds up in tumor cores due to the fast pace of cellular growth outmatching the growth in supply. Hypoxia affects nearly every step in the metastatic cascade which a cancer cell journeys through, but it is likely to affect fibroblasts in the cancer milieu. In this study using a combination of time-resolved control of oxygen dynamics, gene expression analysis, and functional assaying of stromal control of breast cancer invasion, we characterized the effect hypoxia has on breast cancer associated fibroblasts. Furthermore, we also tested if hypoxia induced effect on CAF gene expression is altered when hypoxia is unstable, or fluctuating, as is the case in many regions in hypoxic tumor. Oscillations in hypoxia has recently begun to be described with the advent of new imaging technologies, and we suspect that unstable hypoxia is likely much more common than is perceived. In addition, we have recently described that oscillations in HIF-1 activity can emerge in a subpopulation of hypoxic cancer population with lactate build up in the extracellular space, which can drive HIF-1 degradation through the non-canonical chaperon mediated autophagy. In either cases, both stable, and oscillatory HIF-1 transcriptional activity are physiologically important cues present in the tumor microenvironment, and we know nothing yet about their role on CAFs.

Gene expression analysis broadly revealed that stable hypoxia can cause broad changes in gene expression in CAFs mostly on expected lines. We found that pathways associated with glycolysis, and fibroblast activation were significantly increased in CAFs in response to hypoxia. However, myogenic TFs were predicted to be downregulated, suggesting that while ECM production, ECM-cellular adhesion, and integrin mediated signaling was activated, the actomyosin contractile machinery was possibly not activated, or even inactivated. This surprising finding was confirmed by traction force microscopy, which revealed a significant reduction in force generation by CAFs in hypoxia. Overall, these gene expression changes were accompanied by increased stromal invasibility, a phenotype which we have shown is selected and regulated.

Oscillatory signals are received and processed by the transcriptional machinery likely by “averaging” the amplitude over time, resulting in a sub-maximal dose response. Therefore, the default and trivial assumption for oscillatory hypoxia vs stable hypoxia is of partial reversal in gene expression, which is mostly observed. However, we had previously shown that gene expression in cancer could be specific to oscillatory hypoxia, wherein the patterns did not reflect a sub-hypoxic response. Here, we found that overall, oscillatory hypoxia resulted in partial reversal of nearly all pathways which were upregulated in hypoxia, except for those associated with Actin networking, and capping of pointed end filaments suggesting maturation and stabilization of actin filamentous network. TFM showed that oscillatory hypoxia reversed the trend in hypoxia, with a significant increase in traction force generation, and interestingly, increased variability within the population. Our stromal invasion assay also showed an increase in invasibility by CAFs for MDA-MB-231 invasion. Although our data showed that increased maturation of actomyosin assembly by TGFb (or oscillatory hypoxia), and therefore increased fibroblast contractility was associated with increased invasibility, we posit that more caution is required before we make strong conclusions. Contractile forces, we suggest, act as amplifiers in a basal phenomenon. If cells offer resistance, increased contractile force generation may amplify that response, and vice versa. Force generation within a single cell, homotypic coupling within fibroblasts, and heterotypic coupling between fibroblasts and cancer cells, as well as the fundamental balance between resistance and abetment of invasion are all likely to play a composite role. Future studies should attempt to delineate these composite phenotypes.

Together, our study document the gene expression changes in breast CAFs in hypoxia, as well as those which are specifically affected by fluctuations in environmental hypoxia, and association with a phenotypic effect of increased stromal invasibility to breast cancer. Overall, we show that hypoxia and its fluctuations can alter CAF vulnerability to cancer invasion, likely contributing to cancer dissemination and metastasis.

## Methods

### Isolation of Cancer Associated Fibroblasts

Fresh breast cancer tissue was obtained from Biorepository department at UConn Health Center which was collected from a patient undergoing surgery and placed in sterile RPMI. CAFs were isolated using the ex-plant method. briefly, the tissue was cut into small pieces (approximately 1 mm3) and placed in tissue culture plates containing DMEM/F12 supplemented with 10% FBS and 1% penicillin-streptomycin. The plates were then incubated at 37°C in a humidified incubator with 5% CO2. After 48 hours, the CAFs migrated out of the tissue pieces and adhered to the plates. The tissue pieces were then removed and the CAFs were cultured for further experiments. The CAFs were maintained in DMEM supplemented with 10% FBS, 1% penicillin-streptomycin and 10 ng/ml basic fibroblast growth factor (bFGF).

### Demography Information

Invasive Carcinoma - Histologic Type: Invasive lobular carcinoma **ER+PR+HER2-**

Histologic Type Comments: Invasive lobular carcinoma predominantly shows a solid architectural pattern; tumor cells are negative for E-cadherin. Glandular (Acinar) / Tubular Differentiation Score 3

Nuclear Pleomorphism, Score 2 | Mitotic Rate: Score 1 (<=3 mitoses per mm2) | Overall Grade: Grade 2 (scores of 6 or 7) | Tumor Focality; Single focus of invasive carcinoma

### Bioinformatics Methods

The raw RNAseq reads were aligned to the GRCh38 human genome and transcriptome using hisat2^30^, the SAM file was sorted using samtools^31^, and reads aligning to each coding gene were counted using featurecounts^32^. The read counts were subsequently analyzed using DESeq2^33^ on the R platform to give gene wise log fold change ratios and p-values. The p-values were corrected for multiple testing using the Benjamini-Hochberg method^34^ (i.e., false discovery rates). Gene set enrichment analysis for Gene Ontology (GO)^35^ and KEGG^36^ pathways was performed using the Kolmogorov-Smirnov test in R using the fgsea package^35^. All GO analysis was performed using the biological process ontologies. For all heatmaps, read counts were first transformed to transcripts per million (TPM values), and then z-scores calculated by normalizing each gene independently across the samples shown in each figure. Transcription factor activation scores were calculated using Ingenuity Pathway Analysis from the

### Accelerated Nanopatterned Stromal Invasion Assay (ANSIA)

Cell patterning was performed using a PDMS stencil fabricated using a stereolithographic plastic mold. The stencil was placed on the substrate after washing with isopropanol and drying with N2 steam. The device was kept in a vacuum chamber to remove air bubbles under the stencil. Cancer cells were labeled with CellTracker™ Green CMFDA dye (Thermo Fisher Scientific) following the manufacture’s guideline. Briefly, MDA-DB-231 cells are culture to 90% confluence and washed once with PBS. Then, cells were incubated with 1uM of dye prepared in HBSS or PBS for 20 minutes. Cells were then detached and washed for at least 3 times in 50 ml tubes with PBS. Afterwards, cells were seeded at a density of 5 × 105 cells and attached to the gas permeable plate overnight. The stencil was removed carefully using blunt-end tweezers. The unlabeled stromal cells were seeded at a density of 5 × 105 to fill and attach that area covered by the stencil before. The unattached cells were washed off after 5 h of incubation. The plate was mounted on the microscopy stage. Microscopy was performed on Zeiss Observer Z1 microscope with Colibri.2 LED-based fluorescence and Apotome.2 sectional illumination using a 20× objective.

### Quantification of Extent of Cancer Cells’ invasion into Stroma

Cancer cells labeled with green tracker were imaged at t=0 and t=24 hr. The area occupied by the cancer cells were measured by using Region of Interest (ROI) panel in the Fiji software. ROIs were measured by tracing of boundaries with the freehand tool module in the software. The normalized extent of invasion was calculated by dividing total Area by the length of the initial cancer–stroma interface.

### Traction force microscopy

Traction force gels were fabricated using protocols previously described^37^. Briefly, coverslips for gel attachment were cleaned with ethanol and sonication, treated with air plasma, and activated with 0.5% glutaraldehyde and 0.5% (3-Aminopropyl)triethoxysilane (Sigma Aldrich). Coverslips for beads coating were treated with air plasma and coated with 0.01% poly-L-lysine (Sigma Aldrich) before coated with red carboxylate-modified microspheres with diameters of 0.1 um (Thermo Fisher). Gel precursor solution containing 7.5% acrylamide and 0.15% bis-acrylamide was degassed for 30 minutes and mixed with 0.1% tetramethylethylenediamine and 0.1% ammonium persulfate before sandwiched between silane-activated coverslips and bead-coated coverslips for 20 minutes. After peeling off the bead-coated coverslips, the resulting traction force gels were coated with 30 ug/ml collagen type I using sulfo-SANPAH (Thermo Fisher) overnight at 4 °C. Gels were sterilized under UV for at least 2 hours before cell seeding. Images containing microbeads location before and after cells trypsinization were recorded using Zeiss Observer A1 microscope. Traction forces were calculated following protocols previously described^38^.

## Acknowledgements

We thank our National Cancer Institute R37CA248161 for funding the research presented in the manuscript.

